# Evaluation of Diffusion Tensor Imaging in the Corpus Callosum on a Portable 100 mT MRI System

**DOI:** 10.64898/2026.04.24.717780

**Authors:** Philip Kenneth Lee, Suen Chen, Sijie Zhong, Changyue Wang, Zhiyong Zhang

## Abstract

Low-cost portable MRI has the potential to improve the accessibility of MRI, but new acquisition methods and protocols must be developed and evaluated to accommodate the reduction in SNR and greater impact of system imperfections. Diffusion tensor imaging (DTI) is a candidate tool for monitoring population health, but the bias and variance of quantitative diffusion tensor-derived metrics must be evaluated prior to designing such studies.

DTI of the corpus callosum was performed on an in-house, portable 100 mT MRI system using a slab diffusion weighted Fast Spin Echo with radiofrequency (RF) encoding. Slice coverage was restricted to the corpus callosum to shorten scan time and reduce sensitivity to large rigid motion.

In vivo DTI images were obtained in two healthy volunteers with nominal voxel size 50 mm^3^, scan time 25 minutes, and two different volunteers using nominal voxel size 25 mm^3^, scan time 35 minutes. Mean diffusivity (MD) and fractional anisotropy (FA) coefficients of variation were estimated in the 50 mm^3^ acquisition using a bootstrap approach and compared to resolution-matched data obtained on a conventional 1.5T system. MD / FA maps were compared quantitatively and qualitatively.

Mean MD values in the corpus callosum obtained on the 100 mT system were within 10% of the reference 1.5T acquisition, but FAs were underestimated by 20-30%. The corpus callosum median MD coefficient of variation was 3.7%, and the median FA coefficient of variation was 7.5%. FA maps obtained at 100 mT had an elevated FA noise floor and color FA maps had lower apparent resolution but some white matter tracts were still distinguishable.

**Highlights:**

- Diffusion Tensor Imaging (DTI) of the corpus callosum was performed on a portable 100 mT MRI scanner with 50 mm^3^ voxels in 25 minutes scan time.
- Mean Diffusivity estimates in the corpus callosum obtained at 100 mT and 1.5T differed by less than 0.1 × 10^-3^ mm^2^/s.
- Some white matter tracts were visible in color Fractional Anisotropy maps obtained at 100 mT but FA maps were underestimated by 20– 30% when compared to a resolution-matched 1.5T acquisition, and had lower apparent resolution.

## 1. Introduction

Portable MRI has the potential to increase MRI access to low and middle income countries, remote areas, and point-of-care settings (Wald et al., 2020). These systems are easy to site, can obtain images without an RF shielding room (Marques et al., 2019), and can be powered by a standard electrical outlet. Portable MRI systems eliminate hardware elements commonly found on superconducting systems to reduce cost, weight, and spatial footprint, and operate at lower main magnetic field strengths which reduces SNR (Marques et al., 2019). Longer acquisition times and larger voxel sizes balance the reduction in SNR. The adoption of portable MRI systems represents a tradeoff between overall image quality or scanning efficiency, that is offset by a potential increase in the number of people that can be imaged.

Diffusion Tensor Imaging (DTI) (Basser et al., 1994; Alexander et al., 2007b) has been proposed as a method for monitoring population health (Littlejohns et al., 2020), early detection of Alzheimer’s disease (Acosta-Cabronero and Nestor, 2014; Ruiz-Rizzo et al., 2024), and multiple sclerosis assessment(Pokryszko-Dragan et al., 2018; Kolasa et al., 2019). Fractional Anisotropy (FA) is a biomarker that can be derived from the diffusion tensor, and is sensitive to changes in tissue microstructure (Jellison et al., 2004; Tournier et al., 2011). DTI acquisitions are challenging because of the increase in scan time used to obtain the additional diffusion direction encoding dimension. The diffusion tensor estimate is also sensitive to the SNR and artifact level of individual images. When few diffusion directions are obtained (Skare et al., 2000), diffusion tensor estimates are sensitive to coherent ghosting artifacts or signal dropout in the individual images. Furthermore, trends in FA are only visible at the population level, making it difficult to assess the quality of results based on a small number of participants.

Due to the challenges associated with DTI acquisitions, some tradeoffs must be made. We elected to design a ∼25 minute DTI acquisition that targets the corpus callosum, trading off slice coverage to improve image quality and reduce sensitivity to large rigid motion over long scan times. The corpus callosum has been proposed as an important brain region in the study of cognition and disease progression, and has been studied for its role in aging (Lebel et al., 2010; Pietrasik et al., 2020), tramautic brain injury (TBI) (Kim et al., 2015; Arenth et al., 2014), and autism (Alexander et al., 2007a). Since the corpus callosum is a large white matter tract, it can be segmented even when larger voxels are acquired, and it can fit into a 4-5 cm slab which is ideal for acquisition efficiency and shot-to-shot phase navigation(Miller and Pauly, 2003; Uecker et al., 2009).

Diffusion weighted EPI is the dominant acquisition technique on super-conducting systems, but it is unsuitable on portable MRI systems because the many hardware components that have enabled its success are not available. We recently described an alternative DW acquisition approach using RF-encoded DW-FSE (Lee et al., 2026a), which uses quadratic phase cycling (Le Roux, 2002) to eliminate artifacts caused by non-CPMG magnetization. Non-CPMG magnetization arises from system imperfections and motion-sensitizing diffusion gradients, which must be accounted for in the reconstruction (Lee and Hargreaves, 2022). RF-encoded slabs improve SNR efficiency compared to 2D imaging, and leverage the low SAR advantage provided by lower main magnetic field strengths. A calibration scan is used to map systematic phases incurred by eddy currents and residual fields from different diffusion encodings, which are also incorporated into reconstruction.

The purpose of this work is to characterize the accuracy of DTI-derived metrics acquired on an in-house 100 mT portable MRI system. Brain tractography has previously been demonstrated using DTI on a 64 mT portable system (Gholam et al., 2026), but the accuracy and precision of diffusion tensor-derived metrics was not evaluated. Such an evaluation is important for the design and planning of population-based studies on portable systems Jones and Cercignani (2010); Lauzon and Landman (2013). The accuracy of the reconstruction signal model and possible effects on the diffusion weighted images and DTI metrics were evaluated in phantom experiments. The image SNR and uncertainty of diffusion tensor-derived metrics were evaluated in vivo using a bootstrap approach. Finally, DTI-derived metrics obtained at 100 mT were compared to those acquired on a 1.5T system.

## 2. Methods

### 2.1. Sequence

The multiband DW-FSE sequence is shown in Figure 1. Multiband RF pulses with non-zero DC encodings permit self-navigated imaging, and avoid Gibbs ringing artifacts which occur if Fourier encoding is used in the slice direction with few kz slices. A complex Hadamard matrix with order 10 and the non-zero DC property was obtained in (Best and Kharaghani, 2012) and a permuted version of this matrix was used for slice encoding, (shown in Supporting Information Table T1). Pre-excitation gradients with amplitude equal to the diffusion gradients reduce systematic phase errors due to eddy currents and residual fields. Bipolar phase encoding compensates phase accumulations due to concomitant fields(Lee et al., 2026b). Crushers are negated to avoid mixing with the stimulated echo formed from the area imparted by the excitation slice select gradient.

**Figure 1:**
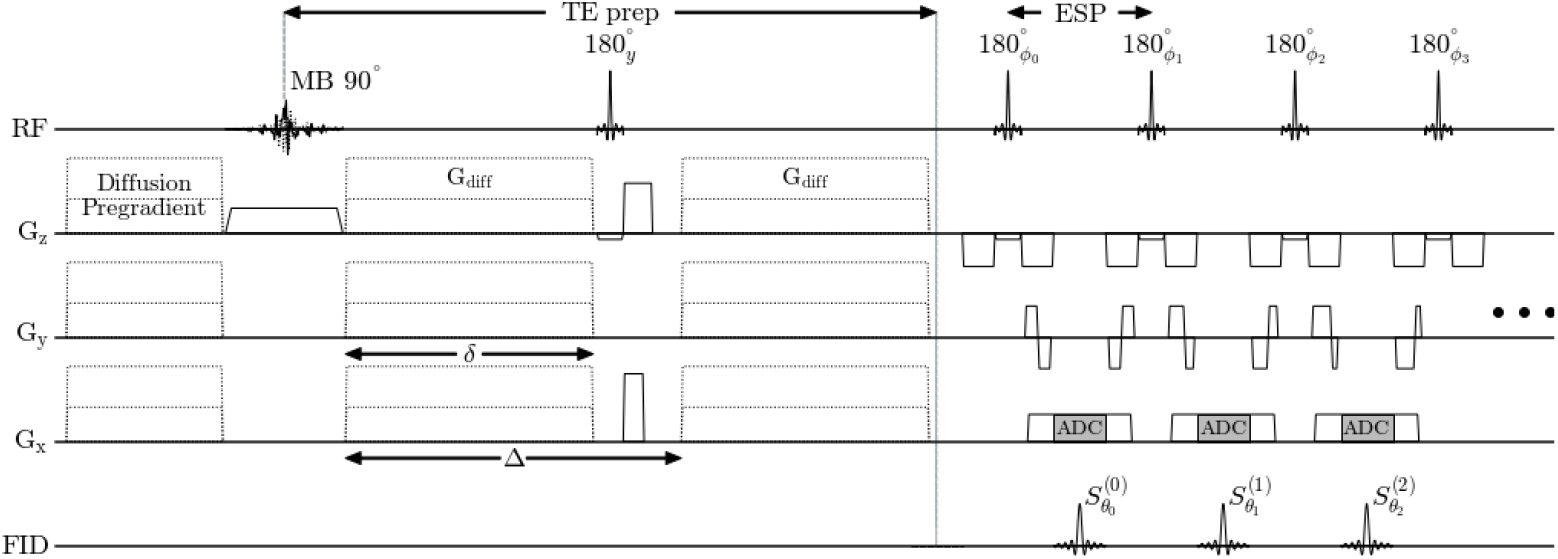
Sequence diagram for diffusion weighted multiband Fast Spin Echo with RF-encoded slabs. The quadratic transmit phase for the *i*th refocusing pulse *ϕ*_*i*_ maintains a stable magnitude signal in the presence of non-CPMG magnetization but must be demodulated with the receive phase *θ*_*i*_ according to the relationship described in Le Roux (2002).

At 100 mT, the T_1_ of white matter (WM) and gray matter (GM) shortens considerably (∼350 ms) when compared to 3T (WM/GM T_1_ ∼900 / 1400 ms, Bojorquez et al. (2017)), but the T2 only increases slightly to ∼90–100 ms (O’Reilly and Webb, 2022) from ∼70 ms at 3T. The T_1_ of CSF is also shortened but to a lesser extent than WM/GM.

The TR that maximizes:

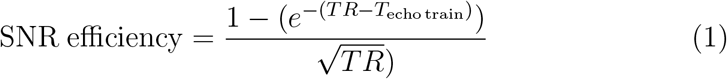

in WM/GM assuming *T*_echo train_ 250 ms is *TR* = 850 ms, with *TR* = 600 ms to *TR* = 1450 ms providing 90% of the maximum SNR efficiency. The short TR induces T_1_ weighting in CSF which is apparent in the b = 0 s/mm^2^ image and reduces partial voluming and Gibb’s ringing artifacts caused by CSF (Chou et al., 2005).

### 2.2. Reconstruction

The linear model differentiates systematic phases ℰ due to eddy currents, concomitant fields, and residual fields from motion-induced phases *P*. This distinction is useful because systematic phases can be measured in a calibration scan and later incorporated into the reconstruction.

Data acquired with the multiband non-CPMG quadratic phase increment refocusing sequence (Le Roux, 2002; Lee et al., 2026a) can be reconstructed by solving the following linear system:

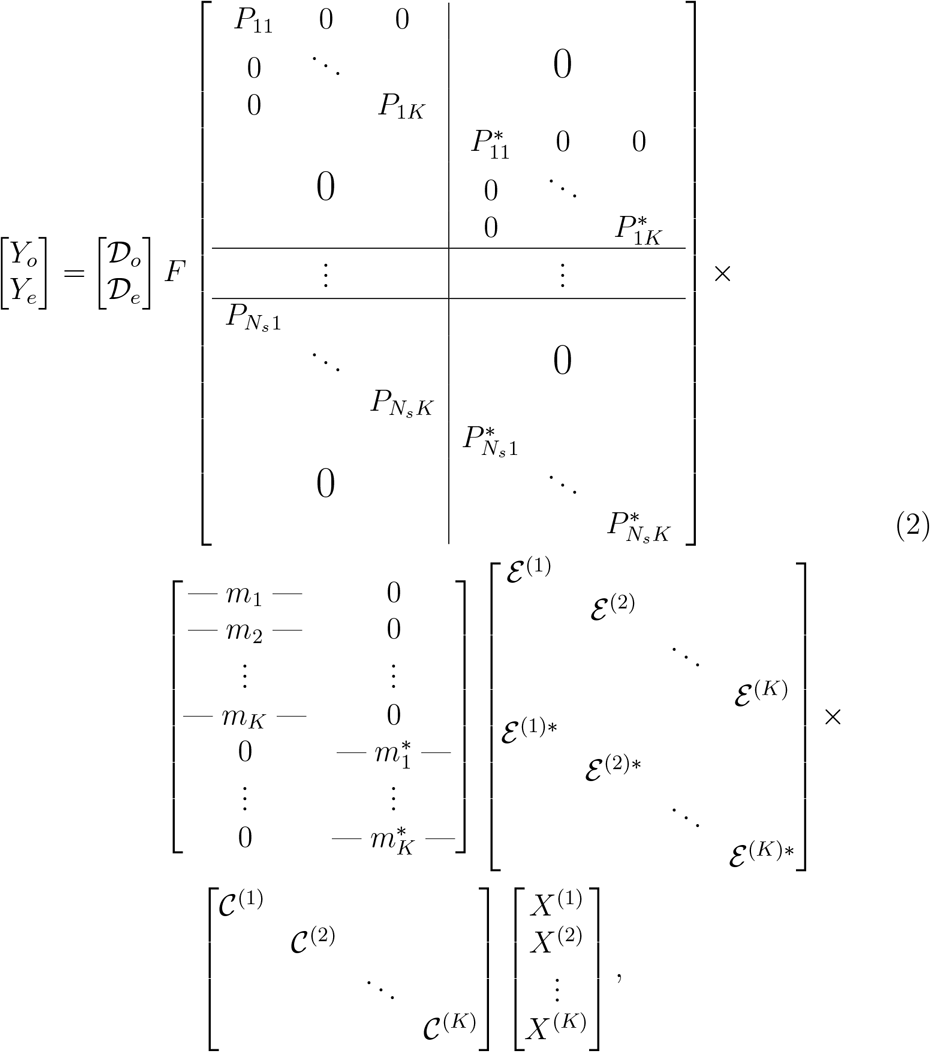

where *K* is the number of RF-encoding bands, *N*_*s*_ is the number of shots, 𝒟_*o*_, 𝒟_*e*_ are block matrices representing the odd and even echo sampling patterns for each shot, *P* is the motion-induced phase, *m*_*k*_ is the RF-encode for RF-encode index *k*, ℰ ^(*k*)^ is the systematic phase error for band *k*, 𝒞 ^(*k*)^ is the CPMG phase reference for band *k*, and *X*^(*k*)^ is the reconstructed image for band *k*.

In the previous implementation of the multiband DW-FSE sequence (Lee et al., 2026a), refocusing pulses were designed to satisfy CPMG for each band with its excitation pulse pair. Relaxing this constraint on the refocusing pulse allows a singleband refocusing pulse to be used. A singleband refocusing pulse has a shorter duration than the multiband refocusing pulse which improves readout efficiency. For example with B_1 max_ ∼ 125 *µ*T, the shortest possible duration 4-band and 10-band TBW 8 refocusing pulses are 2.5 ms and 6 ms respectively, but a singleband refocusing pulse can have a duration as short as 1–1.5 ms. Relaxing the identical phase constraint on the refocusing pulse requires slight adjustments to the linear system used for reconstruction. Compared to Lee et al. (2026a), the non-identical phase refocusing pulse changes the action of RF-encoding (matrix with rows equal to *m*_*k*_ in Equation 2), where the odd echos are measured through the complex conjugation of the RF-encoding matrix to reflect the changes in rotation axis.

The CPMG phase reference(Lee et al., 2026a) 𝒞 is obtained from fully sampled even and odd echo images and allows even and odd echo k-space lines to be reconstructed jointly. Generating virtual odd echo data (Lee and Hargreaves, 2022) allows even and odd echo data to be used to jointly reconstruct *P*. An analogous concept to the CPMG phase reference in DW-FSE is EPI’s use of positive and negative polarity readouts with phase encoding disabled. This calibration allows EPI to use even and odd lines jointly after applying a Nyquist ghost correction with a constant + linear phase model.

To obtain ℰ, a calibration phantom is scanned with diffusion gradients on, and the data is reconstructed using Equation 2 with *P* = *I* (phantom is static), and *X*^(*k*)^ = 1 (phantom assumed to be uniform). Finally, a 3rd order 2D polynomial to the image phase. To obtain *P* for each shot and RF-encode, Equation 2 is solved with *X*^(*k*)^ = 1 and *P* is the variable.

#### 2.3. Phantom Experiment

A custom 20 cm diameter cylindrical phantom filled with peanut oil was imaged using the single-slab DW-FSE sequence with parameters described later in Section 2.4.1. Using peanut oil as a medium is informative because it has an MD ∼ 100× lower than free water (Verma et al., 2017). The signal intensity of peanut oil after diffusion weighting should therefore exhibit no measurable attenuation for b-values commonly acquired in vivo. Signal errors in this experiment can be attributed to either systematic phase errors in the forward model, or system instabilities. This phantom was also used for estimating ℰ.

The phantom data was reconstructed using RF-encoded phase navigators reconstructed from low-resolution data acquired at the end of the echo train. Using the motion phase navigator in the phantom data is not strictly necessary, but its inclusion allows the entire forward model to be evaluated.

### 2.4. in vivo Experiments

Four healthy volunteers were recruited following local IRB approval and informed consent. Different diffusion directions were independently reconstructed using the linear model in Equation 2 and 𝓁1-wavelet regularization applied in 3 dimensions. Next an in-plane Gaussian window was applied with FWHM in k-space at 0.9 k_max_, followed by Marchenko-Pastur Principal Component Analysis (MPPCA) denoising on the magnitude images (Veraart et al., 2016). Diffusion tensors were fit using nonlinear least squares (Koay et al., 2006). The *dipy* implementation of MPPCA denoising and DTI fitting was used (Garyfallidis et al., 2014).

Acquisitions were performed in an RF-shielded room on a 100 mT portable low field scanner with an SmCo_3_ permanent magnet. The system weighs approximately 800 kg, and has dimensions 1.0 × 0.85 × 1.45 m^3^ (width × length × height). The gradient magnitude used for diffusion encoding was 20 mT / m, the maximum employed peak B_1_ was 135 *µ*T, and the maximum applied slew rate was 50 mT / m / ms. Acquisitions were axial with phase encode direction L/R.

The motion phase navigator was estimated from 6 lines acquired at the end of the echo train sampling the k-space center. The nominal phase navigator resolution was 25 × 35 × 40 mm^3^. Multiband-10 excitation pulses with TBW 8, duration 6 ms, and a single band TBW 8, duration 1.5 ms refocusing pulse were designed using SLfRank (Ong et al., 2023).

#### 2.4.1. Experiment 1: Evaluation of Diffusion Tensor Metrics in Corpus Callosum

First, the image SNR, and MD and FA Coefficient of Variation (CoV = *σ/µ*) were evaluated. Image SNR was estimated using the noise estimate provided by MPPCA (Veraart et al., 2016). The MD / FA CoV was obtained in the corpus callosum using a bootstrap approach by retrospectively (Heim et al., 2004) sampling with replacement from an acquisition with additional NEX per direction (termed the data superset) in two healthy volunteers. In the high field images, voxels with MD > 1.25 mm^2^/s were deemed to be contaminated by CSF and were removed from DTI metric comparisons.

The nominal acquired voxel size was 3.5 × 3.5 × 4.0 mm^3^ = 50 mm^3^ with 1 mm slice gap. The DTI acquisition used the following parameters: 20 direction Jones scheme b = 850 s/mm^2^ (3 NEX each) and b = 0 s/mm^2^ (4 NEX) (Jones et al., 1999), diffusion gradient plateau time *δ* = 30 ms, time interval Δ = 40 ms, diffusion pregradient time 30 ms, TE preparation 75 ms, ESP 8.5 ms, TR 800 ms, readout bandwidth 16 kHz, matrix size 64 × 64 × 10, 4 shots per RF-encode, ETL 22 (16 for primary train + 6 for phase navigator).

The total scan time is: 10 RF encodes × 4 shots × (20 × 3 NEX + 1 × 4 NEX) directions × 800 ms TR ≈ 34 minutes. Retrospective reconstruction subsets used 2 NEX per direction to simulate 24 minute acquisitions over 100 bootstrap iterations.

A reference 20-direction DTI was acquired on a 1.5T Siemens Aera system using DW-EPI with 3.5 × 3.5 × 4 mm^3^ voxels, matching the resolution acquired on the portable 100 mT system. Other relevant parameters: b-value = 0 / 900 s/mm^2^ (1 NEX each), TE 80 ms, TR 4000 ms, 20 channel head / neck coil, parallel imaging factor 2, SPAIR fat suppression, scan time 1:30.

#### 2.4.2. Experiment 2: Higher Resolution DTI Acquisition on Portable System

To assess the effects of T2 blur and other systematic errors on apparent resolution, a higher resolution DTI acquisition was performed in two different healthy volunteers and compared to the 1.5T acquisition. The higher resolution acquisition 100 mT had a nominal 2.5 × 2.5 × 4.0 mm^3^ = 25 mm^3^ voxel size. All other parameters were identical to Experiment 1 except: matrix size 100 × 100 × 10, 6 shots per RF-encode, readout bandwidth 32 kHz, 2 NEX per diffusion direction, total scan 35 minutes.

## 3. Results

### 3.1. Phantom Experiment

Images from the oil phantom reconstructed *with* the motion phase navigator are shown in Figure 2, and demonstrate the effect of system instabilities and residual systematic phase errors in the forward model. The distribution of errors is dependent on direction, and the standard deviation of the signal ratio averaged across directions was 5%. If the object is assumed to be uniform intensity, uncorrected phase errors could be considered as a multiplicative, coherent noise since ghosting is proportional to the signalLiu (2016).

**Figure 2:**
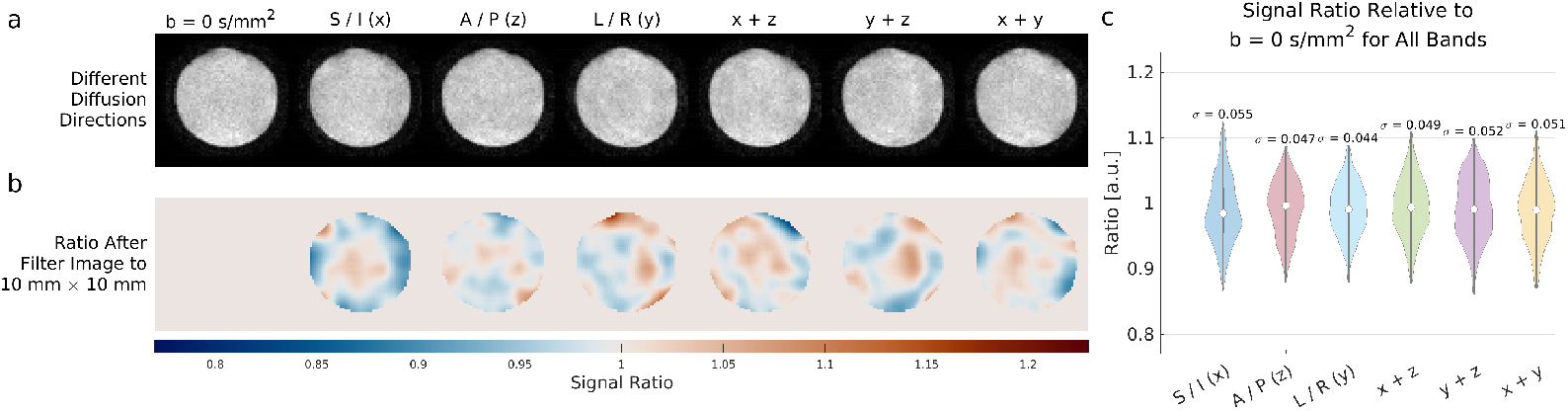
(a) Reconstructed images obtained in oil phantom. The main magnetic field is oriented in the z direction (A/P). (b) Maps of the signal ratio and the (c) distributions per direction for the entire slab. Signal errors due to systematic phase errors are smoothly varying. The mean absolute error (relative to ratio = 1) averaged across all directions is 0.039.

This effect limits the maximum image SNR to ∼ 20–25, and increasing the number of averages will not reduce this noise.

### 3.2. in vivo Experiments

Figure 3 shows diffusion weighted images from Volunteer 1 for select diffusion encoding directions in slices containing the corpus callosum. Image SNR maps as estimated by MPPCA are also shown. The median image SNR in the low field diffusion weighted images was 17.3 in Volunteer 1 and 18.3 in Volunteer 2.

**Figure 3:**
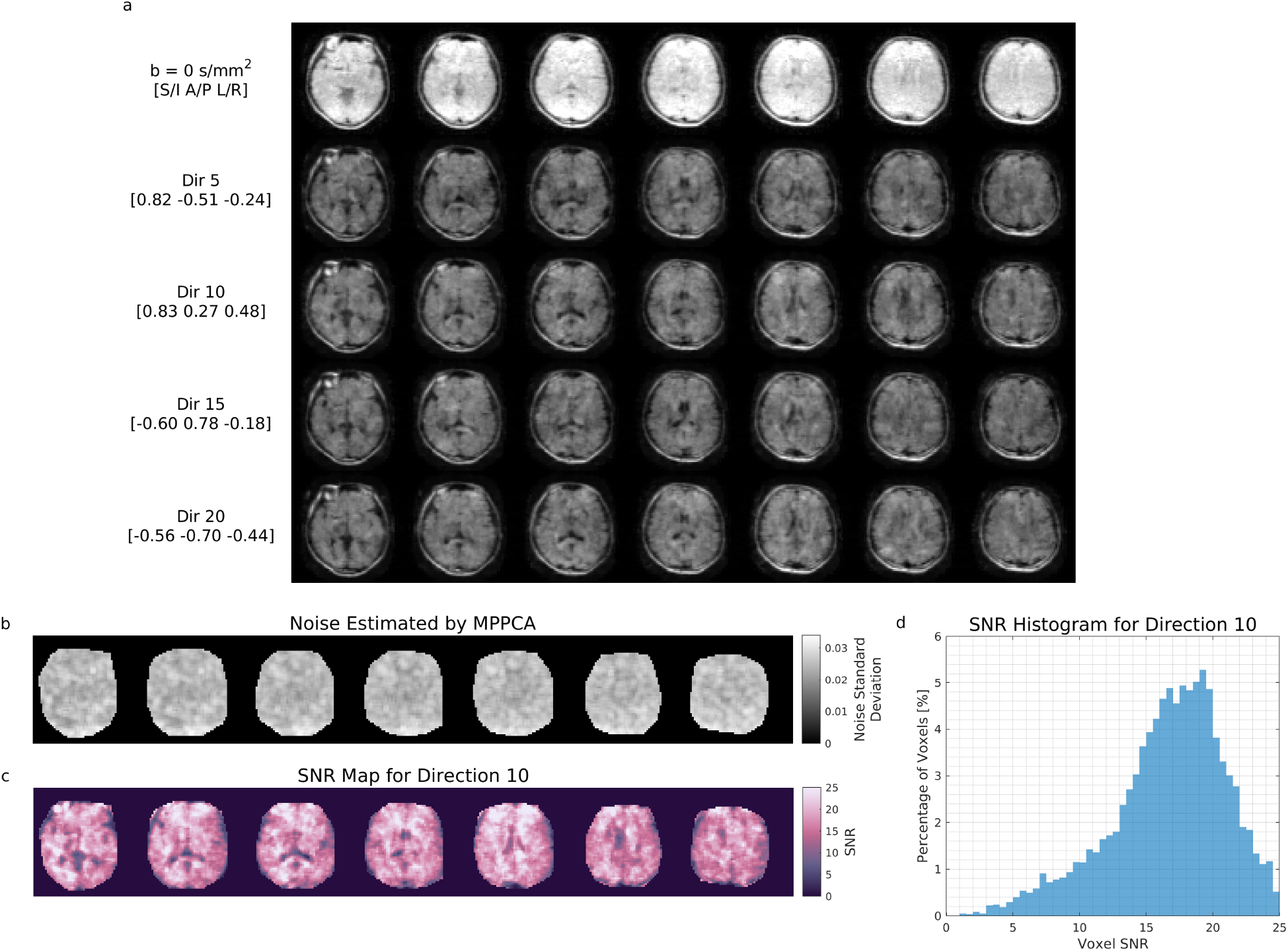
a) Volunteer 1 reconstructed diffusion weighted images after windowing and MP-PCA denoising (voxel volume 50 mm^3^). b) Noise estimates obtained from MPPCA with the (c) computed SNR map and (d) SNR-per-voxel histogram for one diffusion direction.

Figure 4 shows diffusion-tensor derived metrics calculated from Volunteer 2. The MD and FA mean ± standard deviation in the corpus callosum of Volunteer 1 were, 0.80 ± 0.02 *×* 10^−3^ mm^2^/s and 0.51 ± 0.15 respectively. Visible white matter tracts include: the corpus callosum, corona radiata, and cingulum.

**Figure 4:**
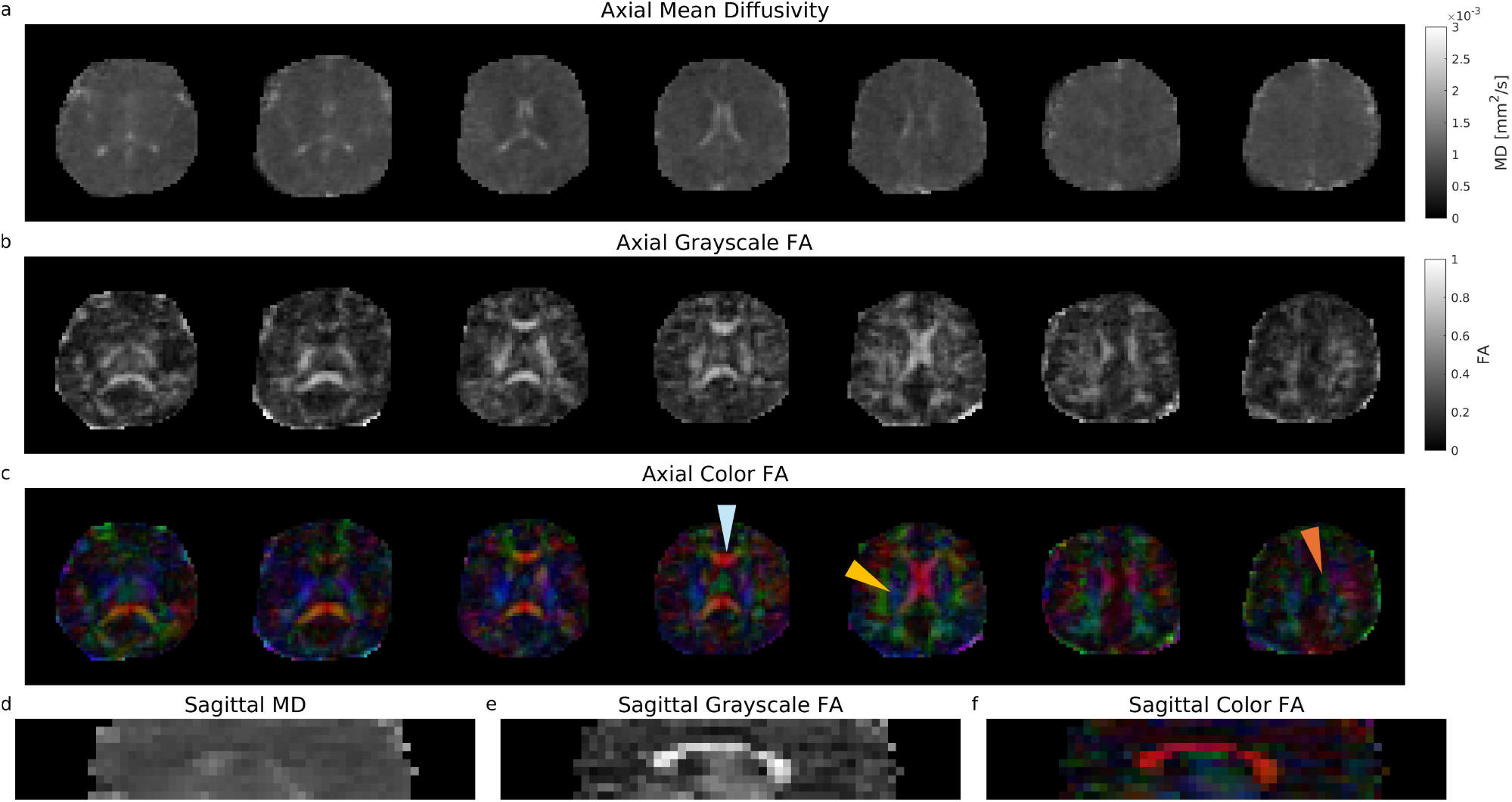
Volunteer 2 images acquired with voxel volume 50 mm^3^. a) MD, b) grayscale, and (c) color FA maps with their sagittal reformat d-f) calculated from images in Figure 3. Visible white matter tracts include: the corpus callosum (light blue arrow), corona radiata (yellow arrow), and cingulum bundle (orange arrow).

Figure 5 compares images from Volunteer 2’s retrospective 25 minute acquisition to those obtained at 1.5T. In Volunteer 2, the low field MD and FA in the corpus callosum were 0.82 ± 0.01 *×* 10^−3^ mm^2^/s and 0.59 ± 0.08. The high field MD and FA were 0.85 ± 0.02 *×* 10^−3^ mm^2^/s and 0.75 ± 0.15. FA is comparatively underestimated in the low field images. The corpus callosum median coefficient of variation obtained using bootstrapping was 3.7% for MD, and 7.5% for FA. Due to the small number of samples in the superset (3 NEX and 20 directions), the coefficient of variation is underestimated by 10–20% Chung et al. (2006).

**Figure 5:**
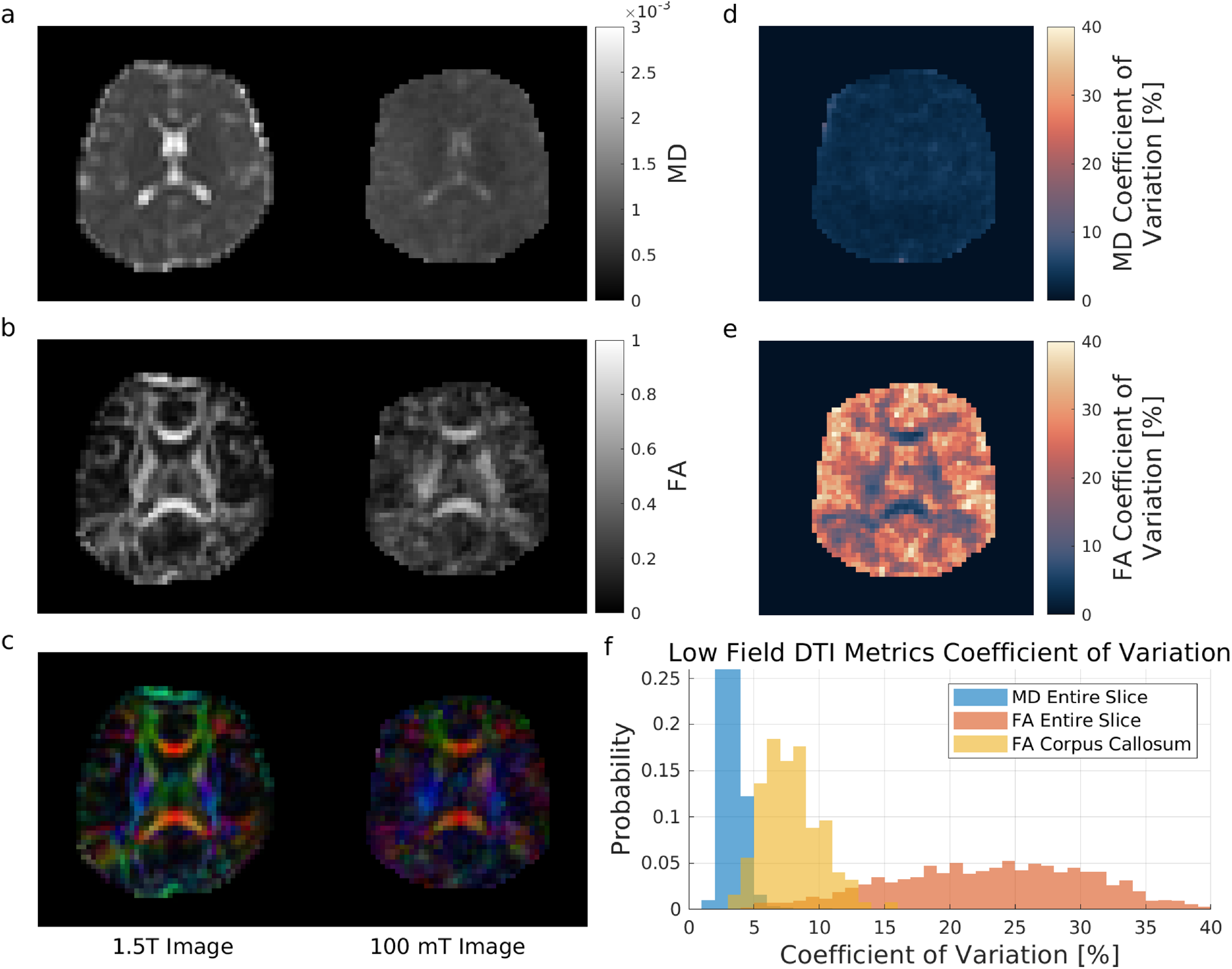
Volunteer 2 image comparison between resolution-matched (voxel volume 50 mm^3^) 1.5T acquisition and retrospective 25 minute acquisition sampled *without* replacement at 100 mT. a) MD, b) FA and c) color FA maps. d-f) 100 mT MD and FA coefficients of variation obtained from retrospective reconstructions sampled *with* replacement.

Despite the same nominal acquisition resolution, smaller structures are not visible in images acquired on the portable system. This is due to the combined effects of low SNR, T2 blurring, rigid motion, and signal discrepancies in DW images from uncorrected systematic phase errors. These effects also incur biases in FA, where the FA background noise floor is raised, and FAs in the corpus callosum are underestimated. CSF in the low field portable images has lower MD compared to the 1.5T images due to low SNR and partial voluming with WM/GM. A slice in the same volunteer offset 1 cm in the superior direction has a more favorable comparison for the low field image in terms of visible structures, shown in Supporting Information S1. This suggests that some of the FA discrepancies may be attributable to uncorrected motion-induced phase.

Figure 6 shows images from Volunteer 3’s 35 minute acquisition at 100 mT with voxel size 25 mm^3^. The median image SNR estimated by MPPCA in this higher resolution acquistion was 8.1. Visually, the apparent resolution of the low field images better match the 1.5T acquisition, but smaller structures are not visible due to low SNR.

**Figure 6:**
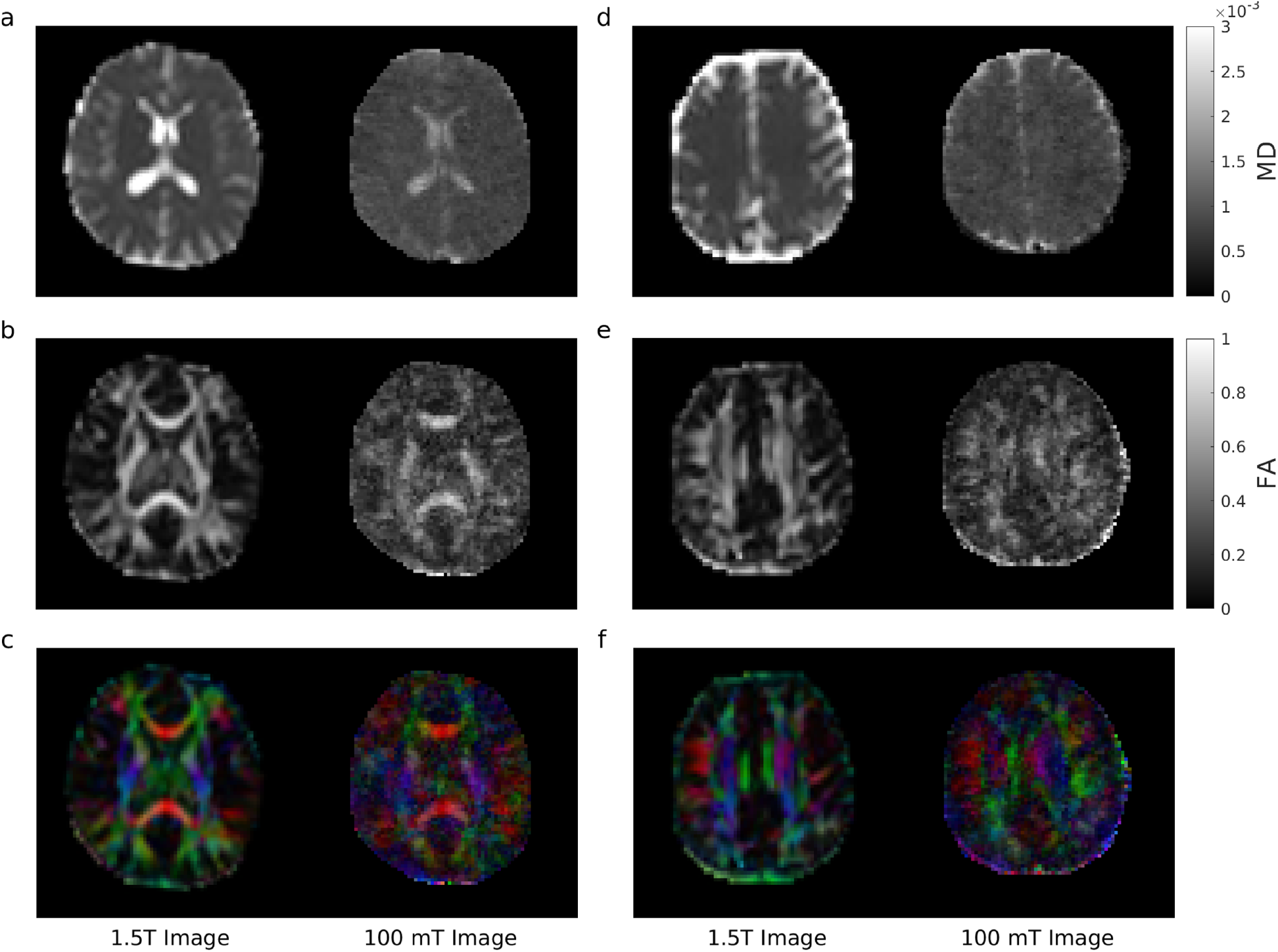
Comparison of images acquired at 1.5T, voxel volume 50 mm^3^, and 100 mT with 35 minutes scan time, voxel volume 25 mm^3^. MD, FA, and color FA maps for a-c) Volunteer 3 and, d-e) Volunteer 4.

Figure 7 compares corpus callosum MD and FA metrics between low field and high field across all four volunteers. Mean MD values at low field obtained in the corpus callosum (ranging from 0.80 – 0.84 × 10^-3^ mm^2^/s across volunteers) were within 10% of the 1.5T acquisition and values in the UK biobank: median mean MD in genu / body / splenium: 0.77 / 0.79 / 0.74 × mm^2^/s 10^-3^ Littlejohns et al. (2020). However, mean FA values in the corpus callosum at low field (ranging from 0.5 – 0.6 across volunteers) were underestimated by 20–30% compared to the 1.5T reference and values in the UK biobank (median across population of mean FA in genu / body / splenium: 0.72 / 0.71 / 0.79).

**Figure 7:**
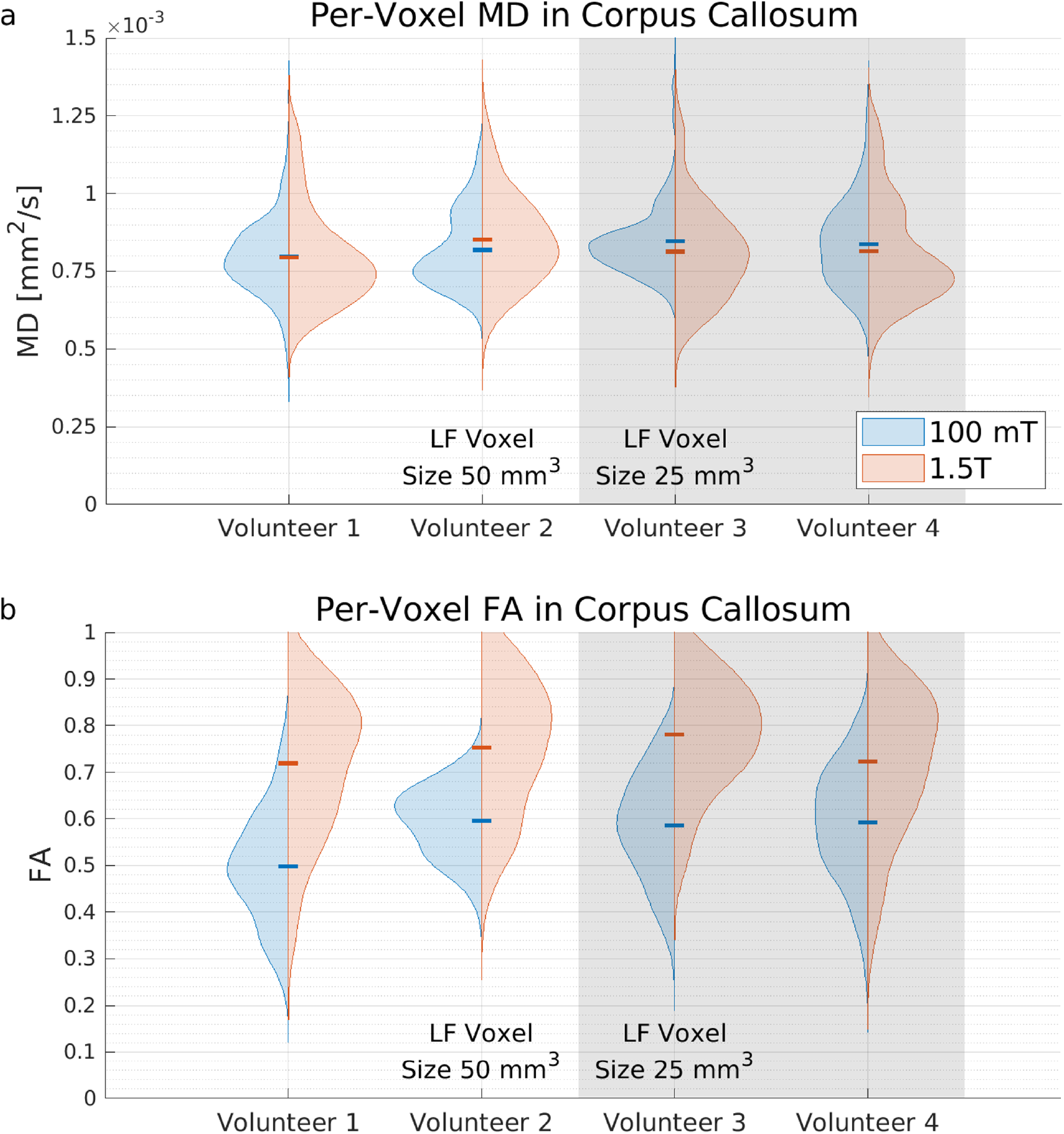
Comparison of a) MD and b) FA values in the corpus callosum obtained at 100 mT and 1.5 T. Mean MD and FA values for each volunteer are denoted with a colored dash.

## 4. Discussion

We have presented a DTI acquisition on a 100 mT portable MRI system which targeted the corpus callosum. DTI metrics were averaged across the entire corpus callosum and compared, but greater number of participants would be required to evaluate differences between the splenium, body, and genu. FAs in the images acquired at 100 mT were comparatively underestimated, and a similar effect was reported in a study at 64 mT (Gholam et al., 2026). Despite FA underestimation on the 100 mT DT images, some white matter tracts were identifiable in color FA maps. Multiple limitations must be considered prior to applying DTI larger population studies.

The first challenge is the low apparent resolution of the acquired images when compared to images with the same nominal resolution obtained on superconducting systems. This is due to the combined effects of: low SNRScheffler (2002), T2 blurring, rigid motion, and signal errors in DW images from uncorrected phases. Care must be taken when comparing image resolution, and such comparisons are most informative when the same hardware and encoding methodology are used, as was done in the 25 versus 50 mm^3^ acquisitions in this work.

Other challenges include those faced by all portable MRI systems, such as removal of electromagnetic interference (EMI) which will reduce the number of usable scans in the study cohort, and the extended scan times making large rigid motions more likely. The integration of motion correction techniques using external sensors Zaitsev et al. (2015) may improve apparent resolution, and cardiac triggering Skare and Andersson (2001) could improve consistency in subjects with higher heart rates.

Furthermore, the design of portable systems varies greatly, and each system design exhibits different artifacts and errors. This study evaluated FAs in one scanner, but inter-scanner repeatability was not evaluated. Due to the greater contributions of system imperfections in portable scanners (Marques et al., 2019), biases between scanners will appear even if they have the same design.

An oil phantom acquisition was used to evaluate errors in the forward model and does not characterize errors due to gradient non linearity which cause deviations from the nominal b-value. Although the systematic error exhibited in the oil phantom appears large with standard deviation 5%, superconducting systems also exhibit scanner-dependent, spurious signal attenuation or enhancement due to gradient non-linearity on the same order of magnitude. Changes in b-value due to gradient non-linearity can be mapped using field probes, or using a phantom with a known MD from 1 – 2 × 10^-3^ s/mm^2^ (Newitt et al., 2015). On superconducting systems over a sphere with radius 10 cm, b-values can vary ±10% of the nominal value (Lee et al., 2020), and this can increase to 15–20% further from isocenter (Newitt et al., 2015). In ice water (MD ∼1.1 × 10^-3^ s/mm^2^), this would cause signal errors ∼5–20%.

### 4.1. DTI SNR Efficiency Comparison Between Diminished Superconducting System and Portable System

One question is how long a DTI acquisition at 100 mT should take, given that many protocols have been developed for DTI at 3T? A simplified argument would be that since the field strength is 30× lower, the SNR is reduced by ∼ 30^3/2^ ≈ 165× (Marques et al., 2019), and the voxel size must increase by that same factor to offset this SNR loss.

This section attempts to improve this estimate by accounting for additional factors including: hardware effects, T2 changes, slice count and sequence considerations, and voxel size. Concretely, suppose that a 3T system such as that used in the Human Connectome Project (HCP) (Uğurbil et al., 2013) were ramped down to 100 mT, and the gradient, shim, active eddy current, and cooling systems were eliminated or replaced with inferior versions.

The HCP DTI protocol acquires: 1.25 mm isotropic voxels, 3 shells: b = 1000, 2000, 3000 s/mm^2^, (90 directions + 6 b = 0 s/mm^2^ = 96) × 2 NEX in 55 minutes, with TE 90 ms. The slice coverage is 111 slices × 1.25 mm = 140 mm. One shell with 140 mm coverage takes 18 minutes, so (20 directions + 1 b = 0 s/mm^2^) requires (20 + 1) / 96 × 18 minutes = 240 seconds = 6 minutes. This 20-direction, 6 minutes protocol will be referred to as *Sample Protocol A*, with an assumed SNR 10 – 15 in the individual images for each direction (Sotiropoulos et al., 2013).

The parameters in the 100 mT portable scanner were: b = 850 s/mm^2^, 50 mm^3^ voxels, 20-direction DTI with 50 mm slice coverage in 25 minutes, or 140 mm in 70 minutes. This 70 minutes protocol with full 140 mm coverage will be referred to as *Sample Protocol B*.

#### 4.1.1. Estimated Effect of Hardware Changes on SNR and Scan Time

The field strength reduction from 3T to 100 mT spans multiple regimes in the dominance of sample noise versus coil noise (Macovski, 1996). Assuming SNR 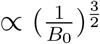 over this range (Marques et al., 2019), the degraded 3T system experiences an SNR penalty 165×, increasing scan time by 165^2^ ≈ **27200**×. The voxel volume acquired at 100 mT was 50 mm^3^ compared to 2 mm^3^ in *Sample Protocol A*. This increases SNR on the degraded 3T system by 25×, or shortens scan time by **625**×.

Increasing the voxel volume reduces the number of slices, which shortens the scan time. Although *Sample Protocol A* uses multiband imaging, the effects on TR will be ignored and we will assume that the reduction in slice count shortens scan time by **3**×.

Diffusion encoding in the Human Connectome Project uses peak amplitude 100 mT/m with slew rate 90 mT / m / ms, and can achieve b-value b = 1000 s/mm^2^ in 30 ms, whereas a gradient system with maximum gradient amplitude 20 mT/m requires 80 ms to reach 1000 s/mm^2^. Since *Sample Protocol A* acquires three shells with EPI and must accommodate the readout train, the TE is 90 ms. Reducing the number of shells for *Sample Protocol A* would reduce TE by ∼ 40 ms, but reducing the maximum gradient amplitude would increase it by ∼ 50 ms. We will assume the net effect leaves the TE unchanged.

Reducing the field strength from 3T to 0.1T increases T2 of WM/GM by approximately 1.5× which creates a 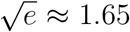 SNR increase, or shortening scan time by **2.7**×.

Superconducting system can employ DW-EPI with high duty cycle due to gradient cooling subsystems, eddy current shielding, and shim coils to image one slice every 125 ms. If DW-EPI were not possible on the degraded system and DW-FSE with longer echo trains had to be used, the time per slice (or diffusion gradient application) would increase to 500 ms. This creates a **4**× time penalty on *Sample Protocol A*.

#### 4.1.2. Final Estimated SNR Efficiency

Starting with *Sample Protocol A*, an acquisition matching scan parameters in *Sample Protocol B* (70 minutes) on the hypothetical diminished system with similar SNR would require approximately 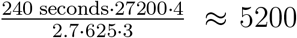 seconds ≈ 85 minutes. Effects that increase acquisition time (such as lower field strength) of the hypothetical diminished 3T system are in the numerator, and effects that decrease acquisition time (such as increasing voxel sizes) are in the denominator. Resolution differences between EPI and FSE due to T2 blurring were ignored.

Based on these estimates, the SNR efficiency of the DTI acquisition on the 100 mT portable system is on the same order of magnitude as the hypothetical *diminished* superconducting system. This estimate is most sensitive to the relationship used between main magnetic field strength and SNR. Due to the lower cost of portable MRI systems per unit compared to superconducting systems, the efficiency of a DTI study may be improved by employing multiple systems in parallel. However, the metric ‘DTI SNR efficiency / monetary unit’ cannot be accurately compare because the cost to manufacture portable systems, and their retail prices are neither public information, nor have they stabilized.

## 5. Conclusion

Diffusion tensor images with restricted slice coverage targeting the corpus callosum can be acquired in a portable MRI system in scan times that may be feasible in larger population studies. Multiband DW-FSE enables this acquisition and exhibits little geometric distortion artifact due to off-resonance. The 100 mT portable system obtained mean MD values in the corpus callosum within 10% of the reference 1.5T acquisition, but FAs obtained at 100 mT were underestimated by 20-30%.

## Acknowledgements

This work was supported by National Key R&D Program of China, 2025YFF0517800; National Natural Science Foundation of China No. 6250010291, No. 62471295; Medical-Engineering Interdisciplinary Research Fund of Shanghai Jiao Tong University YG2025LC01.

## Data Availability

Reconstructed volumes for each volunteer are provided as NIFTI and gif formats in the Supporting Information.

## CRediT authorship contribution statement

**Philip Kenneth Lee:** Conceptualization, Methodology, Software, Validation, Formal Analysis, Investigation, Writing - Original Draft, Writing - Review & Editing, Visualization, Funding Acquisition.

**Suen Chen:** Methodology, Investigation, Resources, Writing - Review & Editing.

**Sijie Zhong:** Methodology, Writing - Review & Editing.

**Changyue Wang:** Methodology, Writing - Review & Editing.

**Zhiyong Zhang:** Conceptualization, Writing - Review & Editing, Funding Acquisition.

