## Supporting Information for "Evaluation of Diffusion Tensor Imaging in the Corpus Callosum on a Portable 100 mT MRI System"

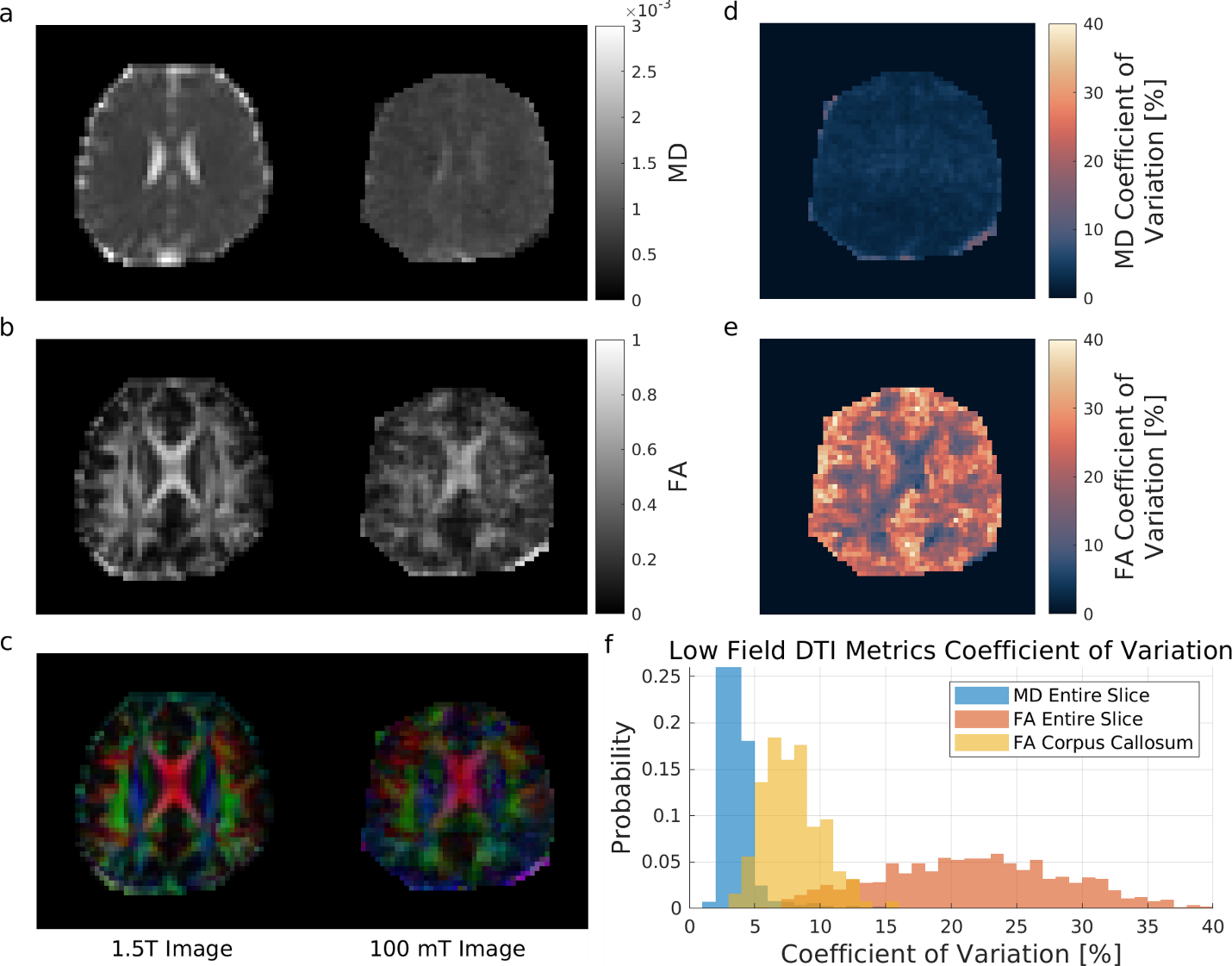

**Supporting Information Figure S1:** Volunteer 2 slice located more 1 cm Superior to slice shown in main text. The color FA maps obtained on the portable low field system have similar structures to the images obtained at 1.5T, but FA values still appear underestimated.

| **RF Encode Index** | **Band**  **1** | **2** | **3** | **4** | **5** | **6** | **7** | **8** | **9** | **10** |
| --- | --- | --- | --- | --- | --- | --- | --- | --- | --- | --- |
| **1** | $\boldsymbol{-i}$ | $\boldsymbol{i}$ | $\boldsymbol{1}$ | $\boldsymbol{-}\boldsymbol{1}$ | $\boldsymbol{-1}$ | $\boldsymbol{1}$ | $\boldsymbol{1}$ | $\boldsymbol{-i}$ | $\boldsymbol{1}$ | $\boldsymbol{1}$ |
| **2** | $\boldsymbol{-i}$ | $\boldsymbol{i}$ | $\boldsymbol{-i}$ | $\boldsymbol{1}$ | $\boldsymbol{1}$ | $\boldsymbol{i}$ | $\boldsymbol{1}$ | $\boldsymbol{1}$ | $\boldsymbol{-1}$ | $\boldsymbol{-i}$ |
| **3** | $\boldsymbol{1}$ | $\boldsymbol{1}$ | $\boldsymbol{1}$ | $\boldsymbol{-i}$ | $\boldsymbol{i}$ | $\boldsymbol{1}$ | $\boldsymbol{-i}$ | $\boldsymbol{i}$ | $\boldsymbol{-}\boldsymbol{1}$ | $\boldsymbol{i}$ |
| **4** | $\boldsymbol{1}$ | $\boldsymbol{1}$ | $\boldsymbol{-1}$ | $\boldsymbol{i}$ | $\boldsymbol{-i}$ | $\boldsymbol{-i}$ | $\boldsymbol{1}$ | $\boldsymbol{1}$ | $\boldsymbol{-i}$ | $\boldsymbol{i}$ |
| **5** | $\boldsymbol{i}$ | $\boldsymbol{1}$ | $\boldsymbol{1}$ | $\boldsymbol{1}$ | $\boldsymbol{-1}$ | $\boldsymbol{-i}$ | $\boldsymbol{i}$ | $\boldsymbol{1}$ | $\boldsymbol{i}$ | $\boldsymbol{-i}$ |
| **6** | $\boldsymbol{i}$ | $\boldsymbol{-}\boldsymbol{1}$ | $\boldsymbol{-1}$ | $\boldsymbol{1}$ | $\boldsymbol{i}$ | $\boldsymbol{1}$ | $\boldsymbol{-i}$ | $\boldsymbol{1}$ | $\boldsymbol{1}$ | $\boldsymbol{1}$ |
| **7** | $\boldsymbol{1}$ | $\boldsymbol{-i}$ | $\boldsymbol{i}$ | $\boldsymbol{i}$ | $\boldsymbol{1}$ | $\boldsymbol{1}$ | $\boldsymbol{1}$ | $\boldsymbol{-1}$ | $\boldsymbol{i}$ | $\boldsymbol{-i}$ |
| **8** | $\boldsymbol{-}\boldsymbol{1}$ | $\boldsymbol{1}$ | $\boldsymbol{i}$ | $\boldsymbol{-i}$ | $\boldsymbol{1}$ | $\boldsymbol{-1}$ | $\boldsymbol{1}$ | $\boldsymbol{i}$ | $\boldsymbol{1}$ | $\boldsymbol{1}$ |
| **9** | $\boldsymbol{1}$ | $\boldsymbol{1}$ | $\boldsymbol{-i}$ | $\boldsymbol{1}$ | $\boldsymbol{-i}$ | $\boldsymbol{i}$ | $\boldsymbol{-1}$ | $\boldsymbol{-1}$ | $\boldsymbol{1}$ | $\boldsymbol{1}$ |
| **10** | $\boldsymbol{-}\boldsymbol{1}$ | $\boldsymbol{-i}$ | $\boldsymbol{1}$ | $\boldsymbol{1}$ | $\boldsymbol{1}$ | $\boldsymbol{1}$ | $\boldsymbol{i}$ | $\boldsymbol{-i}$ | $\boldsymbol{-i}$ | $\boldsymbol{i}$ |

**Supporting Information Table T1:** Complex Hadamard matrix used for RF encoding.
